# Three Dimensional Multi-gene Expression Maps Reveal Cell Fate Changes Associated with Laterality Reversal of Zebrafish Habenula

**DOI:** 10.1101/2020.07.03.182949

**Authors:** Guo-Tzau Wang, He-Yen Pan, Wei-Han Lang, Yuan-Ding Yu, Chang-Huain Hsieh, Yung-Shu Kuan

## Abstract

The conserved bilateral habenular nuclei (HA) in vertebrate diencephalon develop into compartmentalized structures containing neurons derived from different cell lineages. Despite extensive studies demonstrated that zebrafish larval HA display distinct left-right (L-R) asymmetry in gene expression and connectivity, the spatial gene expression domains were mainly obtained from two-dimensional (2D) snapshots of colorimetric RNA *in situ* hybridization staining which could not properly reflect different HA neuronal lineages constructed in three-dimension (3D). Combing the tyramide-based fluorescent mRNA *in situ* hybridization, confocal microscopy and customized imaging processing procedures, we have created spatial distribution maps of four genes for 4 day old zebrafish and in sibling fish whose L-R asymmetry was spontaneously reversed. 3D volumetric analyses showed that ratios of *cpd2, lov, ron* and *nrp1a* expression in L-R reversed HA were reversed according to the parapineal positions. However, the quantitative changes of gene expression in reversed larval brains do not mirror the gene expression level in the obverse larval brains. There were a total 87.78% increase of *lov*^*+*^*nrp1a*^*+*^ and a total 12.45% decrease of *lov*^*+*^*ron*^*+*^ double-positive neurons when the L-R asymmetry of HA was reversed. Thus, our volumetric analyses of the 3D maps indicate that changes of HA neuronal cell fates are associated with the reversal of HA laterality. These changes likely account for the behavior changes associated with HA laterality alterations.

## INTRODUCTION

Along the developmental processes upon forming a functional brain in vertebrates, many differentiating neurons will gather into structurally distinguishable yet functionally coherent or connected clusters of neurons (nuclei or nodes) that contain different cellular characteristics. Proper generation of these compartmentalized and specialized neuronal organizations are needed for facilitating the processing of complex input and output activities within each vertebrate brain (Chatterjee and Li, 2012; Hashimoto and Mikoshiba, 2003; Sugihara and Fujita, 2013).

The highly conserved habenula nuclei (HA) are paired neuronal clusters located flanking to the pineal gland in diencephalon and belong to one of the conserved forebrain to midbrain conduction systems in vertebrate brains (Beretta et al., 2012; Bianco and Wilson, 2009; Roberson and Halpern, 2018; Sutherland, 1982). Studies utilizing rodents have indicated that two obvious compartments called medial and lateral HA are distinguishable in HA, and the medial HA contains about 5 subnuclear groups and the lateral HA contains about 10 subnuclear groups (Aizawa et al., 2012; Andres et al., 1999; Klemm, 2004; Sutherland, 1982). This complex composition of HA is a direct reflection of the diverse physiological functions such as feeding, mating, sleep, diet, stress response and aversion or reward processing it involves (Fore et al., 2018; Klemm, 2004; Sandyk, 1991). Comparing to our understanding in the structure, the neurochemical composition and the physiological function of HA, our current knowledge of HA neuronal generation and specification in vertebrate development is relatively limited.

Studies with mice showed that transcription factors Gbx2, Brn3a, Tcf7l2, Foxg1 and Otx2 each involves in different aspects of HA developments such as HA neurogenesis (Gbx2, Foxg1), HA neuronal proliferation (Gbx2, Otx2), migration (Otx2), differentiation (Tcfl2), and HA neuronal wiring (Brn3a, Tcfl2) (Lee et al., 2017; Liu et al., 2018; Mallika et al., 2015; Quina et al., 2009; RuiZ-Reig et al., 2019). Besides mouse model, zebrafish is also a vertebrate model to study the complex neuronal differentiating and clustering processes of HA because of the presence of plethora of gene markers (Amo et al., 2010). Despite that how each different signaling molecules guide the generation and clustering of different HA neuronal precursors into functionally connected structures are still largely unknown, a picture emerged from several reports indicated that Nodal, Wnt, Notch, Fgf and Shh signaling or ER membrane protein each plays unique role in the early generation (Wnt, Notch, Fgf, Shh, sec61al1) and patterning of HA (Wnt, Nodal, Fgf) (Aizawa et al., 2007; Carl et al., 2007; Concha et al., 2000; Doll et al., 2011; Kuan et al., 2015; Regan et al., 2009; Roberson and Halpern, 2017; Roussigne et al., 2009).

In addition to serve as a model to understand how neuronal clusters are developed in the vertebrate brain, the uncovering of asymmetries in gene expression and connectivity in zebrafish larval HA has promoted zebrafish to become an important model to understand how laterality in vertebrate brains is generated. A number of studies revealed that asymmetric Nodal activation by Wnt signaling combing with Fgf signaling guides the unilateral migration of parapineal (Pp) cells that ensures the establishment of asymmetric gene expressions in left and right (L-R) habenulae (Carl et al., 2007; Concha et al., 2000; Concha et al., 2003; Gamse et al., 2003; Regan et al., 2009). Directions of the asymmetric expressions of *lov, ron* and *nrp1a* in HA could be altered in accordance with the Pp position. Higher expressions of *lov* and *nrp1a* are always ipsilateral whereas higher expressions of *ron* is always contralateral to the Pp position. Further investigation on the correlation between HA laterality and behavioral consequences indicated that establishing HA asymmetry plays important roles in visual and motor behaviors, and in stress and fear responses of zebrafish (Agetsuma et al., 2010; Barth et al., 2005; Dadda et al., 2010; Dreosti et al., 2014; Facchin et al., 2009; Facchin et al., 2015; Okamoto et al., 2012). However, despite that L-R reversals do not necessarily represent a neuroanatomical mirror image, the molecular and cellular basis for the different behaviors mediated by HA with different laterality has not been addressed before (Facchin et al., 2015). In order to understand the differentiation of distinct neuronal lineages within each L-R habenulae in obverse (Pp on left side) or reversed (Pp on right side) zebrafish larval brains, spatial descriptions of gene expression patterns in 3 dimensions (3D) should be adopted to better represent the distribution of different HA neurons because these neurons are organized in 3D in each larvae.

Here we report the generation of 3D gene expression volume maps of four frequently used HA markers (*cpd2, lov, ron* and *nrp1a*) in zebrafish larval brains with different laterality utilizing fluorescent mRNA *in situ* hybridization (FISH), confocal microscopy and customized imaging processing procedures (Gamse et al., 2005; Gamse et al., 2003; Kuan et al., 2007). Through volumetric analyses of the gene expressions, we found that despite the directions of L-R gene expression volume ratios of *cpd2, lov, ron* and *nrp1a* reversed accordingly with the reversal of the Pp positions in larva at 4 day post fertilization (4 dpf), the ratio numbers per se for each genes except *cpd2* in reversed larval brains are not mirroring to the obverse larval brains. Further volumetric analyses of the multi-gene maps indicated that there is a 87.78% increase of total *lov*-*nrp1a* double-positive domains and a 12.45% decrease of total *lov-ron* double-positive domains in HA of L-R reversed larvae at 4 dpf. These observations indicated that cell fate changes are associated with the alterations of HA laterality, and provided an explanation to why L-R reversals do not necessarily represent a neuroanatomical mirror image. These results also indicate that 3D gene expression maps can reveal spatial information of gene expression in brain tissues that are unable to be revealed by 2D image analyses.

## MATERIALS AND METHODS

### Fish Maintenance and Sample Collection

AB strain (wildtype) zebrafish were purchased from Taiwan Zebrafish Core Facility in Academic Sinica. Fishes were maintained and bred according to standard procedures described on ZFIN (http://zfin.org). All larvae were collected from natural spawning and reared/staged in 0.003% PTU, fixed in 4% paraformaldehyde (PFA, Sigma P6148) in PBS overnight at 4°C at 4 dpf then stored in 100% MeOH at –20°C until processed for RNA *in situ* hybridization. Larvae with reversed HA laterality were collected from spontaneously occurred siblings (∼3%) in AB larva (Gamse et al., 2003). All animal procedures were performed in accordance with the protocol approved by the Institutional Animal Care and Use Committee (IACUC) at Academia Sinica (Protocol#11-05-182).

### Fluorescent mRNA *In situ* Hybridization

Digoxigenin (DIG)-labeled *cpd2* RNA probes or fluorescein (FLU)-labeled *lov, nrp1a* or *ron* RNA probes were generated according to previously described procedures (Gamse et al., 2003). The two-color whole mount fluorescent RNA *in situ* hybridization (FISH) was carried out as previously described by Ma et. al. with minor modifications (Ma and Jiang, 2007). Basically, all steps are the same as the chromogen-based *in situ* hybridization (CISH) described by Gamse et. al (Gamse et al., 2003) except that 1 to 600 dilution of anti-DIG-HRP conjugated and anti-FLU-HRP conjugated antibodies (Roche 11207733910 and 11426346910, respectively) in 2% blocking solution (Roche 11096176001) were used for RNA probe detection. For probe detection and presentation, we followed the protocol provided by the commercial company except that all samples were stained in 1 to 80 dilution of Cy3-conjugated or FLU-conjugated Tyrimide substrates for 60 minutes (Perkin Elmer NEL753001 Kit).

### Image Acquisition

The FISH labeled samples were incubated overnight in 50%, then 90% glycerol/PFA before imaging. Fluorescent samples were imaged with Leica TCS SP5 confocal system (Leica GmbH, Germany) using a 10x air objective with 0.3 N.A. Images were scanned with 4X zooming factor and recorded as 512 × 512 pixels and Z-step of 1.98 µm for 24 steps (final 25 frames) at 100 Hz. The resulted voxel size of our images is 0.76 × 0.76 × 1.98 µm^3^. For presentation, Z-projection images were first obtained using “3D projection” function in Leica’s LAS AF software then cropped to proper size using Photoshop CS5 (Adobe Inc., USA).

### Image Process and Visualization

Volumes (Vol.) of each gene expression domains were calculated by combining basic image fragmentation tools (labeling LHA and RHA domains into 2 different materials) and MaterialStatistics module (exporting voxel counts and vol. in µm^3^). Intensity threshold values (0-255) adopted for each gene were 20 for *cpd2* and *lov*, 40 for *nrp1a* and *ron* throughout the vol. calculation and model visualization steps (Supplementary Figure 1A-D’). Signals from each stack were taken once only to avoid the bleaching effects that were obviously affecting the vol. calculation for the signals emitted from FLU-conjugated Tyrimides (green channel) (Supplementary Figure 1E-J). Volumes of overlapped regions were obtained by utilizing Arthmetic and CastField modules to label the overlapped regions first and then were followed by the vol. calculation steps described above. To obtain the Correlation Factors (CFs) between each registering stack pairs, rigid image registrations were performed by utilizing AffineRegistration module with Ouasi Newton as the Optimizer (initial step = 0.1; final step = 0.01) and Correlation as the Register Metric. Data transformation (rigid) was applied with ApplyTransform module using the Lanczos interpolator, then the correlation matrix of the resampled data was computed to obtain the CF utilizing CorrelationPlot module. The stack showing the highest averages of CFs was selected to serve as the reference stack for the subsequent volume averaging and merging steps utilizing AverageVolumes and Merge modules, respectively. Maximum numbers of lattice were preset in Merge module in order to fit the sample size. Resulted new image data from averaging and merging steps were exported out as new data objects that preserved the original coordinate and voxel size. For virtual 3D image visualization, Isosurface module was adopted for each newly merged and averaged data object of each gene with user defined color (grey for C, green for L, blue for N, and yellow for R in this study). Compactify and downsample options with X=2, Y=2 and Z=2 were adopted for generating the final Isosurface views. Images exhibiting different viewing angles of the virtual 3D models were exported from viewer window by utilizing the Snapshot function placed in Amira then they were cropped to proper size using Photoshop CS5 (Adobe Inc., USA).

### Statistical Analysis

Statistics for volume average and standard deviation (SD) and chart generation were performed utilizing Prism 7 software (Graphpad, CA, USA). Unpaired Student’s t-test (two tailed) was utilized to determine significance at the *P* value that is smaller than 0.05. Confidence for *P* value (*p*) is denoted by * equals to *p* < 0.05 and ** equals to *p* < 0.01.

## RESULTS

### Steps for Image Acquisition, Processing and Visualization

To understand the 3D spatial distributions of different neuronal populations in developing HA at 4 dpf, four endogenously expressed and frequently used genes in zebrafish larval HA, the *cpd2, lov, nrp1a* and *ron*, were selected to perform the fluorescent RNA *in situ* hybridization (FISH) (Gamse et al., 2005; Gamse et al., 2003; Kuan et al., 2007; Ma and Jiang, 2007). Each sample was labeled with *cpd2* (the reference gene in this study, named C) plus any 2^nd^ gene of interest (*lov, nrp1a* or *ron* in this study, named L, N or R). The *cpd2* gene was selected to be the reference gene that served as an anchor for combining the expression volume map of other genes because its expression domains were shown to occupy large areas in both left and right (L-R) habenula and its expression was unaffected after removal of parapineal (Pp) cells (Gamse et al., 2003). After successfully obtained 2-color FISH samples, serial images along the dorsal to ventral axis (Z axis) of HA were collected using Leica TCS SP5 confocal microscope. Subsequently, these Z-stacks were subjected to the image processing procedures that utilized Amira software to execute all the image registration, alignment, volume merging and averaging steps (Amira 5.6, FEI, USA; ChingYeh Corp., LTD, Taiwan) (Figure 1A).

**Figure 1.**
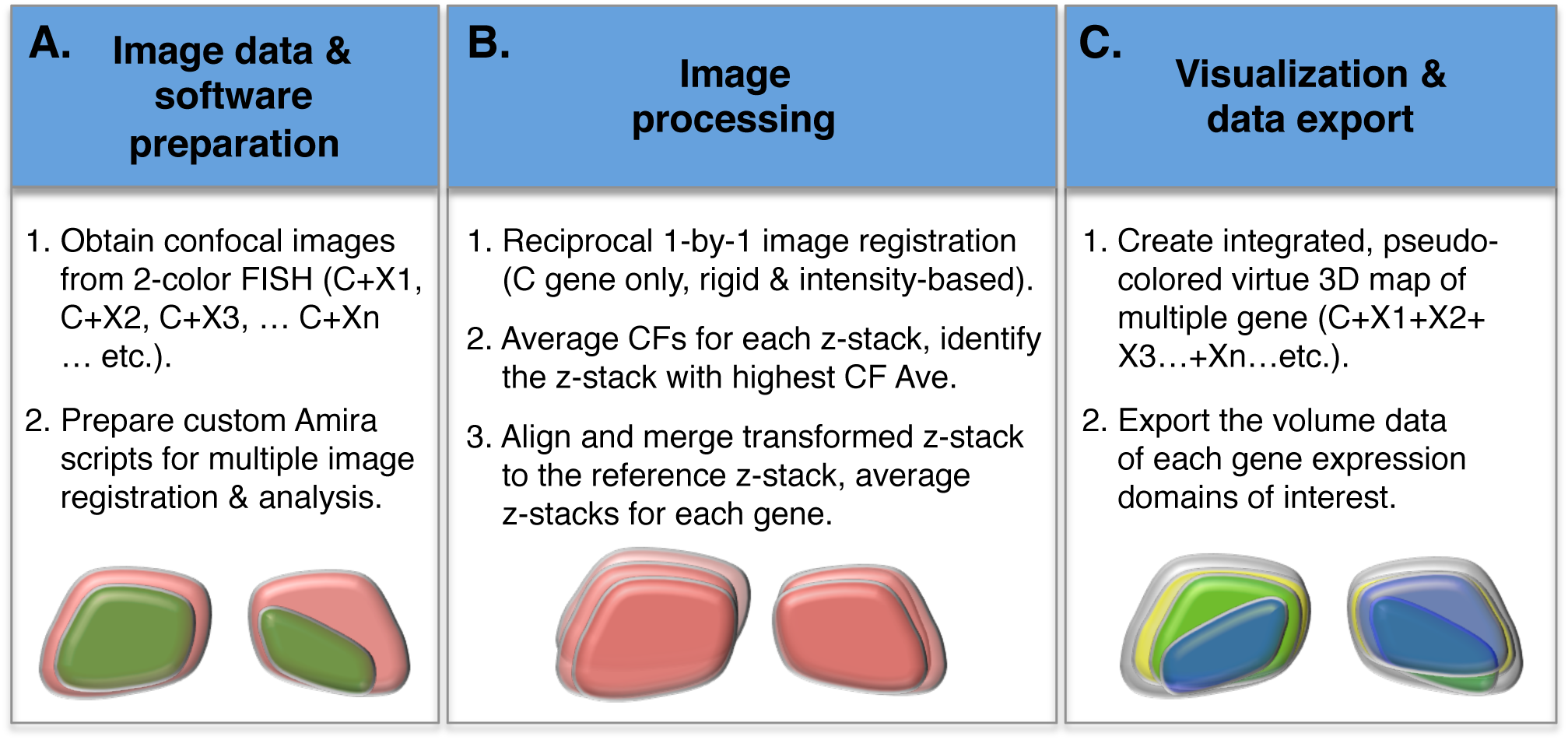
Steps for image acquisition, processing and visualization. (A) Image data acquisition and software preparation. Samples of 2-color fluorescent *in situ* mRNA hybridization (FISH) using reference gene (*cpd2* in this study, named C) plus any 2^nd^ gene of interest (named X) were imaged using confocal microscope. Amira script combining image import, registration, alignment and volume averaging steps was generated for image processing. (B) Perform reciprocal one-to-one registration to obtain correlation factors (CFs) among the C gene channels in each Z-stacks. Compute the averages of the CFs for each stacks. Align all the stacks using the stack showing the highest CF average as the reference stack then merge individual Z-stacks and create averaged Z-stacks as new image objects for each genes. (C) Use the new image objects of each genes to create isosurface rendered, pseudo-colored virtue 3D map of multiple genes for visualization purpose. Compute the volume of each individual Z-stack and export the results in a CVS formatted file.

In order to understand the similarity between each Z-stacks and to identify the Z-stack that best represent the *cpd2* (C gene) sample population, our image processing was initiated by performing reciprocal one-to-one registration to obtain correlation factors (CFs, representing similarities) among the C gene channels of all Z-stacks. The one Z-stack of C gene that generated the highest average of CFs (the highest similarity, Supplementary Table 2A-B) was selected to serve as the reference stack for the subsequent image processing steps. All images that were transformed during the CF calculation steps were subjected to the final image alignment procedures utilizing the reference Z-stack selected from the CF analyses. Volume aligned and averaged virtue Z-stacks for C, L, N and R genes were generated as four new image objects in Amira, and subsequently these image objects were used in visualization and expression domain overlapping analyses (Figure 1B, details in Materials and Methods).

For visualization purpose, we utilized the functional module ‘Isosurface’ in Amira to perform image object rendering. Each rendered virtue 3D image was pseudo-color coded to create the final 3D multiple gene expression maps (see Materials and Methods). The thresholded volumes of each merged stacks obtained during image averaging step could be exported into HxSpreadSheet in Amira as an option for further examination and analysis (Figure 1C).

### Gene *cpd2* Is a Competent Reference Marker for Image Registrations

In order to compare how similar two *cpd2* Z-stacks might be before we start to build the 3D volume map, we performed reciprocal one-to-one image registration as mentioned above (Figure 1B). Utilizing the “Correlation” registering metric in Amira program (details in Materials and methods), all the one-to-one registrations between each Z-stack themselves were resulted in CFs that equal to 1 (Figure 2A, 2E and Supplementary Table 1A-B). These results indicate that our registration steps and metric selections were correctly performed as we expected. In comparison, the CFs from one-to-one C gene registrations between each other Z-stacks in the obverse (Pp on the left) or reversed (Pp on the right) groups would distribute between 0.90 to 0.58 (n=23) or 0.89 to 0.36 (n=14), respectively (Figure 2B-D, 2F-H and Supplementary Table 1A-B). Subsequently we calculated the averages of the CFs for each Z-stacks and found that stack RC4 of the obverse group and stack rLC2 of the reversed group demonstrated the highest averages of CFs (Figure 2I and Supplementary Table 1A-B), indicating that these two stacks would best represent the Z-stacks from each group. Heatmap analysis was adopted to compare the similarities of C gene images between each individual staining subgroups (C+L, C+N and C+R gene in obverse or reversed groups). As can be seen in Figure 2J, the distributions of CF averages were not biased toward any particular staining subgroups in obverse or reversed groups, indicating that the FISH staining efficiencies were similar among the different staining subgroups.

**Figure 2.**
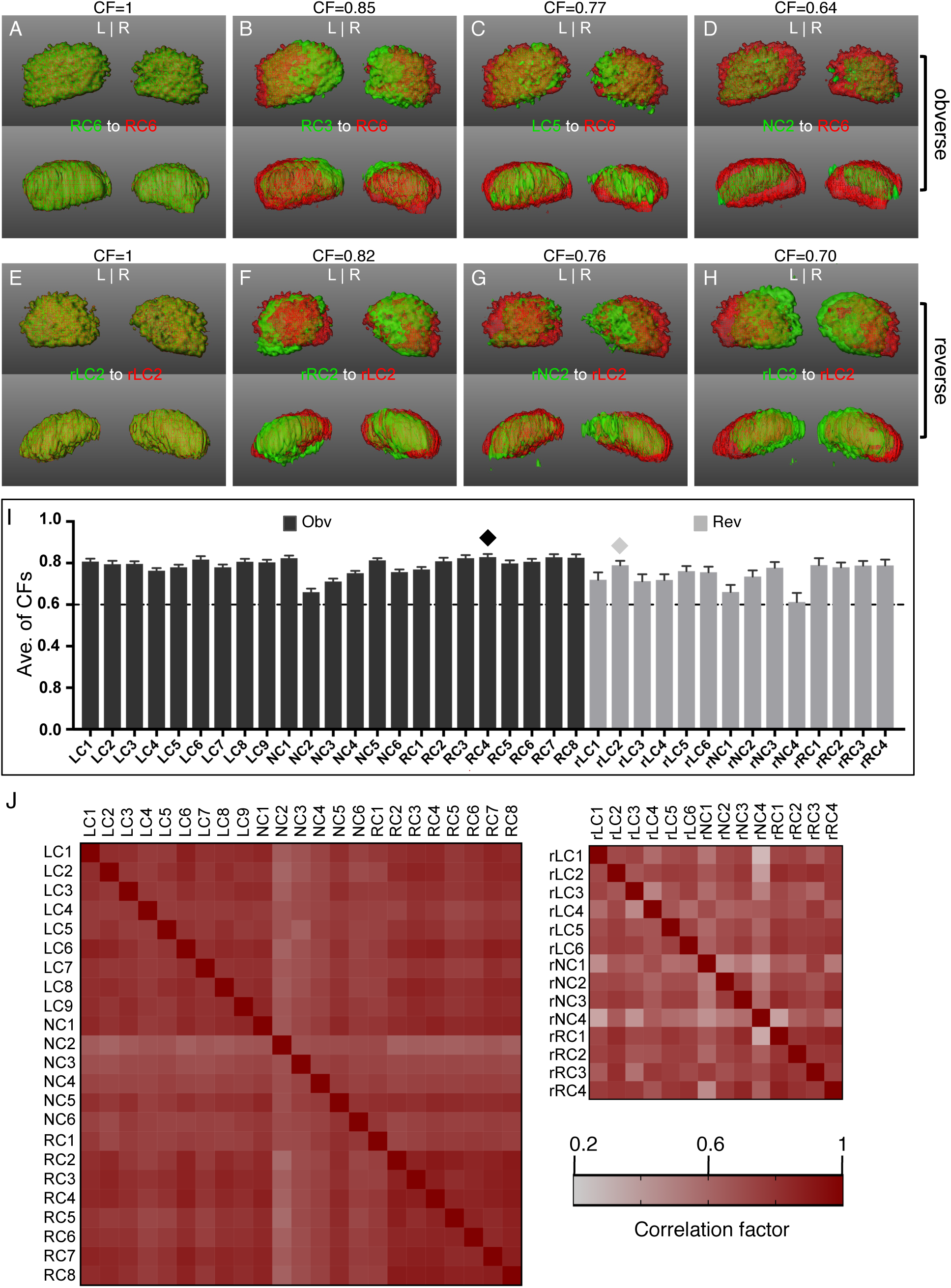
Correlations of *cpd2* expression domains in the habenular nuclei between each image stacks. (A-H) Representative snapshots from image registration analyses for the *cpd2* expression domains. Registration between stacks RC6 against itself (first one in red and second one in green) (A), RC3 against RC6 (B), LC5 against RC6 (C), NC2 against RC6 (D), rLC2 against itself (E), rRC2 against rLC2 (F), rNC2 against rLC2 (G), and rLC3 against rLC2 (H). Top panel for dorsal and bottom panel for anterior views. Correlation factor (CF) represents the fitness between the two registered stacks (refer to the materials and methods). CF equals to 1 representing the highest correlation between the 2 registered Z-stacks. (I) Statistical chart of averaged CFs from each Z-stacks of *cpd2* gene after one-to-one registration. Diamonds mark the selected reference Z-stacks in obverse (Obv) or reverse (Rev) groups. (J) Cluster analyses (heatmaps) of the CFs from one-to-one registration between the Z-stacks of *cpd2* gene. Left or the top-right panels are analysis results for obverse or reverse groups, respectively. Bottom-right panel is the color codes for the heatmaps (also see Supplementary Table 1).

### Gene Expression Volumetric Analyses Revealed Novel Aspects That Are Not Reflected by Colorimetric Fashion of Gene Expression Analyses

Previously, several studies have shown that the asymmetric expressions of *lov, ron* and *nrp1a* in developing HA could be altered in accordance with the position of the Pp, whereas the expression of *cpd2* was unaffected despite of the reversal of the HA asymmetry (Concha et al., 2003; Gamse et al., 2003; Gamse et al., 2005; Kuan et al., 2007; Regan et al., 2009; Roussigne et al., 2009). However, in those studies, the extent of the HA asymmetry has never been addressed at the 3 dimensional gene expression domain level. In our FISH labeling studies, the asymmetric expressions of *cpd2, lov, nrp1a* and *ron* genes in 2D Z-projection images were seemed to be superimposable between the obverse or reversed HA as to what had been reported before (Figure 3A-H) (Concha et al., 2003; Gamse et al., 2005; Gamse et al., 2003; Kuan et al., 2007; Regan et al., 2009; Roussigne et al., 2009). When we examined the expression volumes of each gene in LHA or RHA, we found that the sizes of *cpd2* expression domain were switched accordingly to the position of Pp as expected (99,365±28,282 µm^**3**^ in LHA and 73,728±21,108 µm^**3**^ in RHA of obverse larva, 82,783±16,429 µm^**3**^ in LHA and 93,843±21,546 µm^**3**^ in RHA of reversed larva) (Figure 3I, M and Supplementary Table 2). In addition, the average volumes of *cpd2* expressing domains were relatively larger among the four selected genes in L or R habenula with only one exceptional case in the laterality reversed larva brains. In which case, the averaged *lov* expression domain was larger than the *cpd2* domain in the RHA in (Figure 3I-J, Supplementary Table 2). These observations therefore substantiated that selecting *cpd2* to be the reference gene for constructing the 3D multi-gene expression map of 4 dpf larval HA is a legitimate decision. In comparison, the sizes of the *lov, nrp1a* or *ron* expression domains in L-R reversed HA were however not simply a mirror reversal of their obverse counterparts (78,910±25,974 µm^**3**^ in LHA and 16,141±16,901 µm^**3**^ in RHA for *lov*, 17,782±11,311 µm^**3**^ in LHA and 69,419±20,092 µm^**3**^ in RHA for *nrp1a*, 16,855±10,317 µm^**3**^ in LHA and 26,763±14,701 µm^**3**^ in RHA for *ron* for the obverse group; 50,523±28,535 µm^**3**^ in LHA and 105,076±21,513 µm^**3**^ in RHA for *lov*, 33,578±16,944 µm^**3**^ in LHA and 360±884 µm^**3**^ in RHA for *nrp1a*, 24,737±10,720 µm^**3**^ in LHA and 10,316±4,784 µm^**3**^ in RHA for *ron* for the reversed group) (Figure 3J-M and Supplementary Table 2). This kind of L-R differences has never been reported or addressed before. Therefore, our results demonstrated that 2D image analyses would probably overlook the differences in HA nuclei that were structured in 3D in larval brains. The non-mirroring switch of *lov, nrp1a* and *ron* expressions suggested that there are cell fate changes in reversed larval HA at both the Pp and Pp opposite sides. In addition, the non-mirroring switch of *nrp1a* expression in reversed larvae also suggests that the connectivity is not a mirror reversal to their obverse counterparts, because the efferent axons of HA neurons require Nrp1a to recognize their targets in the IPN (Kuan et al., 2007).

**Figure 3.**
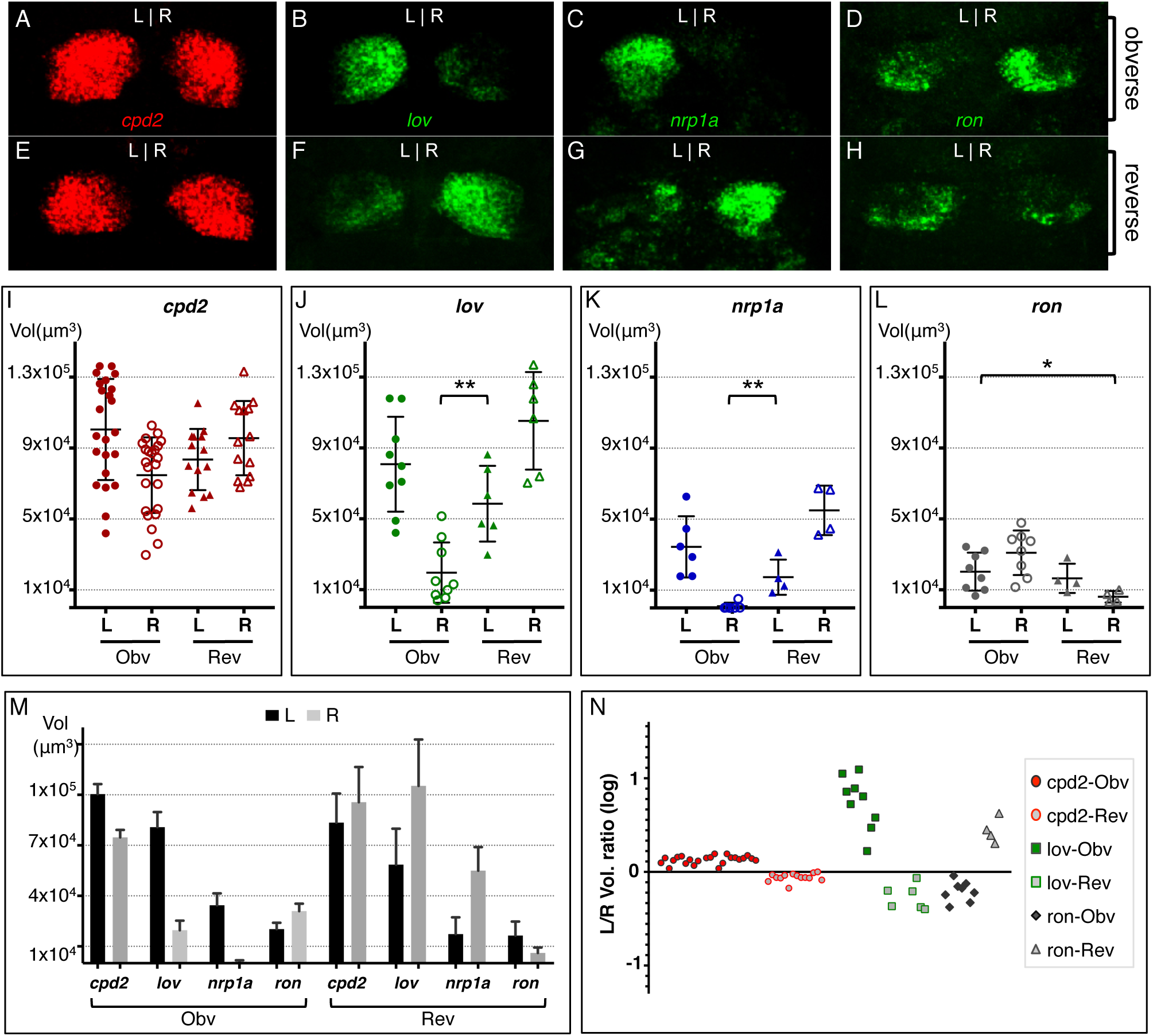
Gene expression volume ratios do not mirroringly reversed with the laterality changes of the habenular nuclei. (A-H) Representative Z-projection images from *cpd2* (A, E), *lov* (B, F), *nrp1a* (C, G) or *ron* (D, H) riboprobe staining in habenula nuclei (HA) of 4 dpf larvae. (I-L) Gene expression domain volumes for each tested riboprobes in left (L) or right (R) HA of obverse (Obv) or reversed (Rev) larval brain (also see Supplementary Table 2). (M-N) Statistical charts showing the averaged gene expression volumes (M) or the L-R gene expression volume ratios (N) from obverse or reversed larval brains. Confidence for *P* value (*p*) is denoted by * which equals to *p* < 0.05 and ** which equals to *p* < 0.01. Only the *P* value indicating a significant change was marked. *P* is 0.0017 in J; is 0.0027 in K; is 0.0299 in L.

### Volume Ratios of The *lov* or *nrp1a* Expression Domains Between Left and Right Habenula Is Significantly Altered in Laterality Reversed Larval Brains

To further understand the relationship of gene expression volumes in each habenulae within the whole sample population, we analyzed the volume ratios of *cpd2, lov* and *ron* between their expression domains in LHA and RHA. The results indicated that the average of *cpd2* expression volumes in LHA is about 1.4 times greater than the expression volumes in RHA (135.9±13.1%), and no sample showed an L to R (L/R) volume ratio lower than 109% in the obverse group (Supplementary Table 2). Reversed yet similar volume ratios were observed in laterality reversed larval samples (L/R is 89.28±9.98%, R/L is 113.36±3.11%.) (Figure 3M-N, Supplementary Table 2). On the other hand, despite the volumes of *lov* expression domains in the HA at Pp side are constantly greater than the volumes in the HA at Pp opposite side in obverse or reversed groups as expected (1,967±933% and 48±29%, respectively), the volume ratios nevertheless are not mirroringly reversed when the Pp positions switched to the opposite sides (Figure 3M-N and Supplementary Table 2). These observations thus demonstrated again that 3D volume analysis could reflect gene expression features that would not be reflected by 2D image analysis.

### Multi-gene 3D Volume Maps for Habenula Nuclei of 4 dpf Larvae

To demonstrate the average expression volumes for each *cpd2* (C), *lov* (L), *nrp1a* (N) or *ron* (R) genes in 4 dpf larval HA, representative snapshots from four different angles (dorsal, ventral, anterior, posterior) of the virtual 3D HA gene expression maps were shown in Figure 4 and 5 for the obverse or reversed groups, respectively. In these figures, the averaged expression domains of *cpd2* gene (in grey) occupied the largest areas in both LHA and RHA in the virtual 3D maps for both obverse and reversed groups (Figure 4A-D, 5A-D). The L, N or R genes each occupied sub-domains in HA that either was overlapped with C gene only or with C and the other genes together to form a complex mixture of cell lineages that can not be clearly seen in the snapshots from traditional 2-color colorimetric staining (Figure 4E-T, 5E-T). For instance, the cell lineages that are CLN gene-positive or CLR gene-positive now can be identified in the 4-gene 3D virtual maps in Figure 4 and 5. Building the multi-gene 3D expression volume map therefore can demonstrate a variety of cell types in the developing HA that can not be shown properly by colorimetric staining.

**Figure 4.**
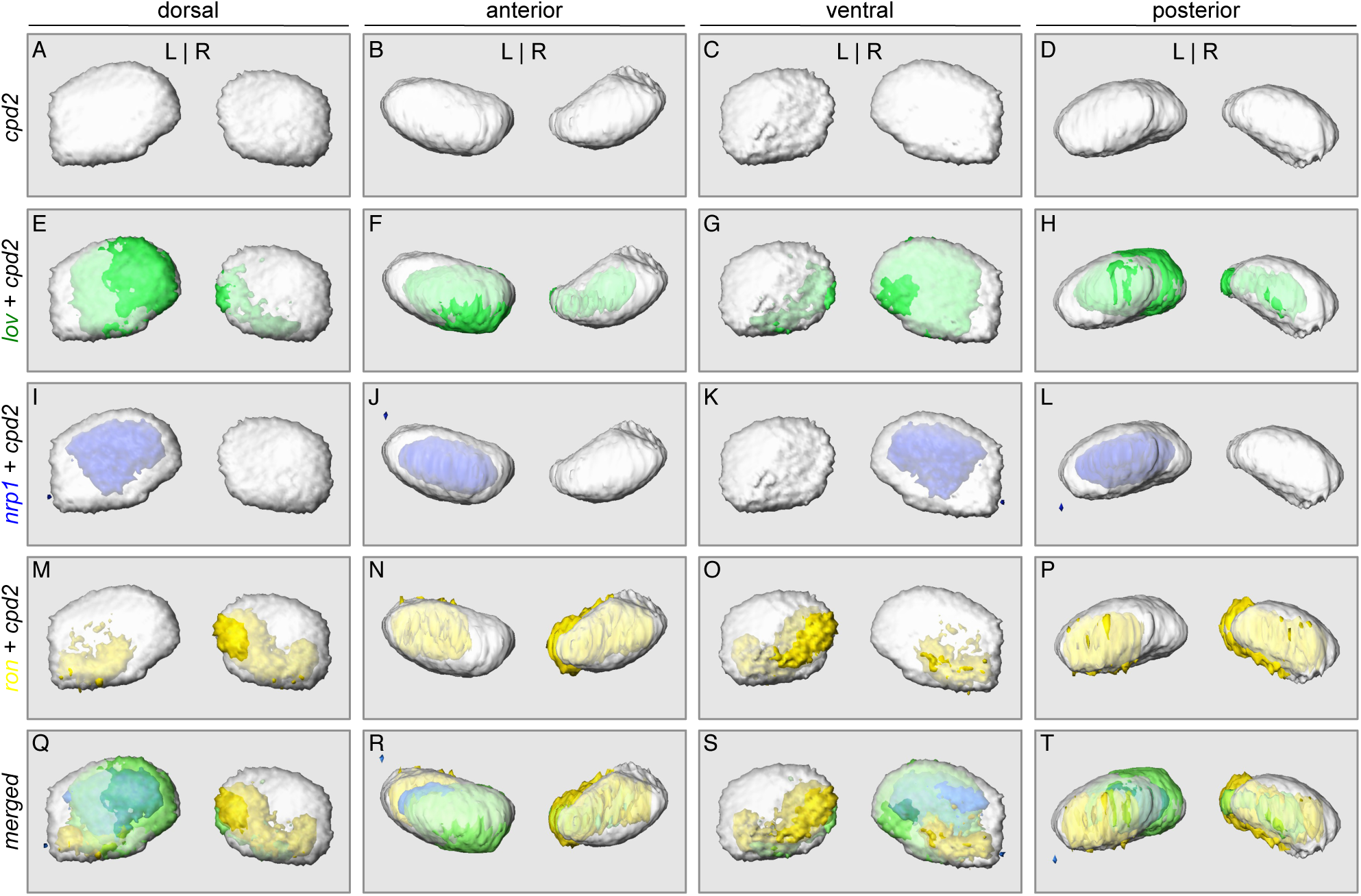
3D multi-gene expression volume map for obverse habenula nuclei of 4 dpf larvae. (A-D) Representative snapshots from four different angles (labeled on top) from the virtual 3D map generated after aligning and averaging 24 Z-stacks of *cpd2* mRNA staining (in gray). (E-P) Representative snapshots from virtual 3D map showing aligned and averaged Z-stacks of *lov* gene (E to H, in green), *nrp1a* gene (I to L, in blue) gene, or *ron* gene (M to P, in yellow) in addition to the averaged virtual images of *cpd2* gene. (Q-T) Snapshots from four different angles from the merged virtual 3D map of all four genes.

**Figure 5.**
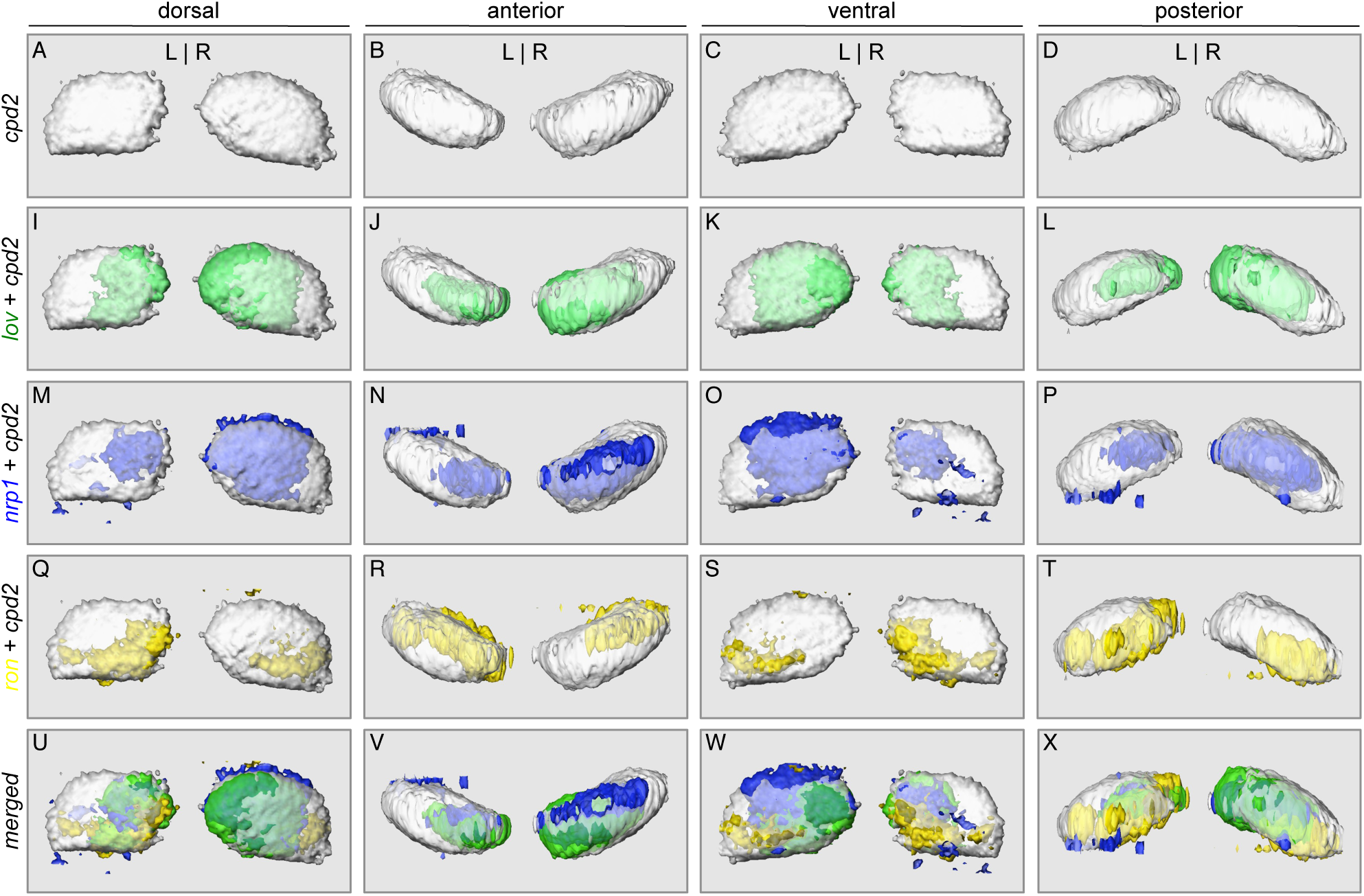
3D multi-gene expression volume map for reversed habenula nuclei of 4 dpf larvae. (A-D) Representative snapshots from four different angles (labeled on top) from the virtual 3D map generated after aligning and averaging 24 Z-stacks of *cpd2* mRNA staining (in gray). (E-P) Representative snapshots from virtual 3D map showing aligned and averaged Z-stacks of *lov* gene (E to H, in green), *nrp1a* gene (I to L, in blue), or *ron* gene (M to P, in yellow) in addition to the averaged virtual images of *cpd2* gene. (Q-T) Snapshots from 4-gene merged virtual 3D images.

### Neuronal Lineage Distributions Are Altered in Laterality Altered Larval Brains

Previously, studies had shown that HA laterality switch in zebrafish brain is associated with the altered neurological functions of HA (Agetsuma et al., 2010; Barth et al., 2005; Dadda et al., 2010; Facchin et al., 2009; Facchin et al., 2015). However, despite that reversal of HA L-R asymmetry does not necessarily represent a neuroanatomical mirror image has been proposed previously, the underlying mechanisms of these behavior consequences are still largely unknown. For instance, whether the neuronal numbers or lineages are altered accordingly in laterality reversed fish has not been explored before. Utilizing the multi-gene 3D maps of 4 dpf HA generated in this study, we further analyze the lineage distributions of *lov* plus *nrp1a* (*lov*-*nrp1a*) double-positive neurons and *lov* plus *ron* (*lov*-*ron*) double-positive neurons in the integrated maps built from obverse or reversed samples. The results indicated that there is a total 87.78% increase of *lov*-*nrp1a* double-positive neurons in HA of L-R revered larvae (Figure 6A-R). Specifically, there is a 53.63% increase of *lov*-*nrp1a* double-positive neurons in habenulae on the Pp side, and an increase from 0 to 12150 µm^3^ of *lov*-*nrp1a* double-positive neurons in habenulae on the Pp opposite side. On the other hand, there is a total 12.45% decrease of *lov*-*ron* double-positive neurons in HA of L-R revered larvae (Figure 7A-R). Specifically, there is a 60.79% decrease of *lov*-*ron* double-positive neurons in Pp-side habenulae, and a 43.45% increase of *lov*-*ron* double-positive neurons in Pp-opposite-side habenulae. These cell fate alterations not only provided us the cellular mechanism that underlies the previously observed behavior changes upon alterations of brain laterality, these results also strongly support our original notion that gene expression volume analysis would reflect the differences that could not be demonstrated by the 2D colorimetric gene expression analysis.

**Figure 6.**
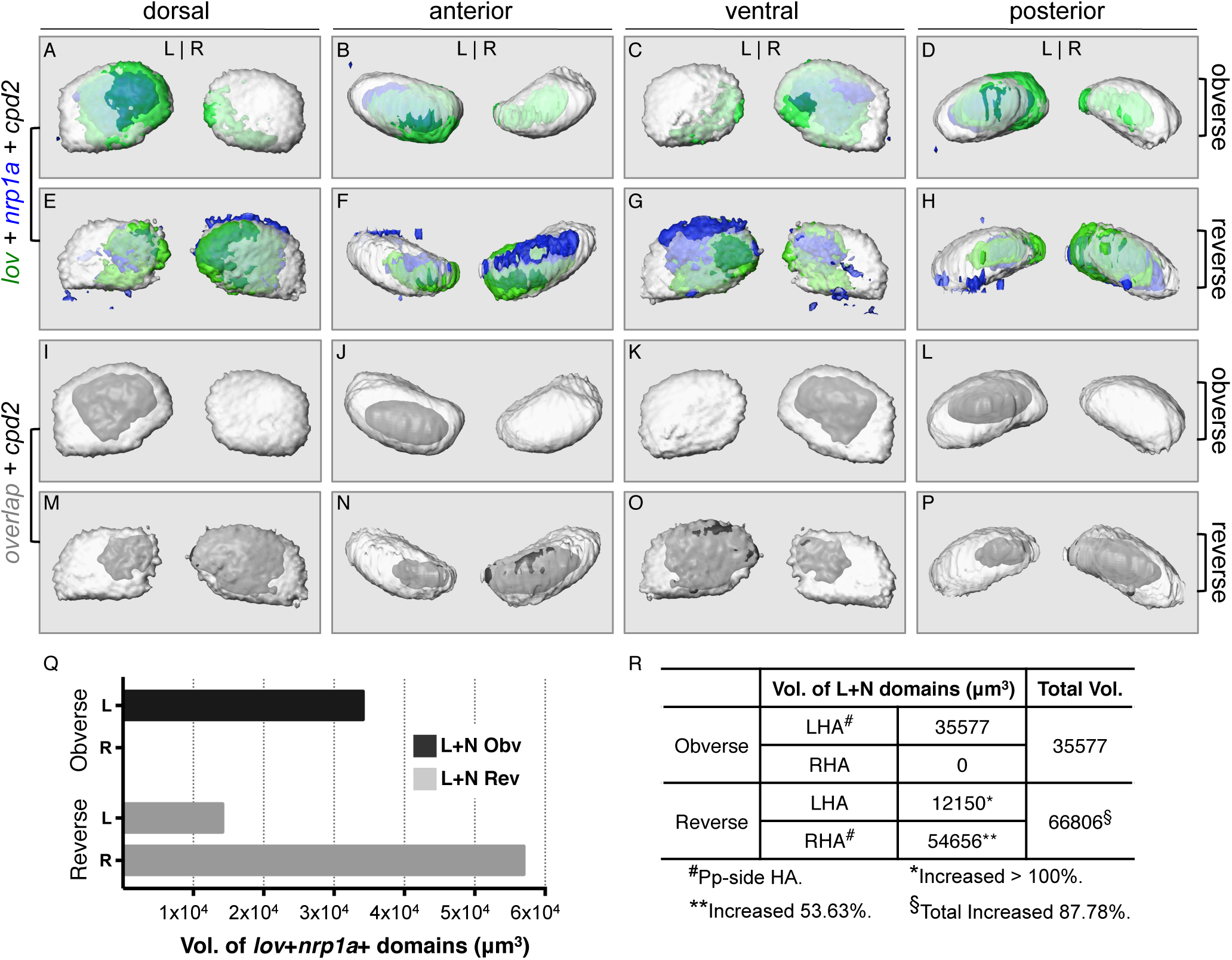
Increase of *lov* and *nrp1a* double-positive domains in left-right reversed habenular nuclei in 4 dpf larvae. (A-H) Snapshots from virtual 3D map from four different angles showing merged maps of *cpd2, lov* and *nrp1a* transcripts from obverse (A-D) or reverse (E-H) groups. (I-P) Snapshots showing *lov* and *nrp1a* overlapped domains (in gray) from obverse (I-L) or reverse (M-P) groups. (Q) Statistical chart showing the expression volumes of the *lov* and *nrp1a* overlapped (L+N) domains in obverse (Obv) or reverse (Rev) model maps. (R). A table showing the computed volumes of L+N domains in the virtual 3D maps. Total changes in percentage was calculated by using (Rev total – Obv total) / Obv total X 100. Same logic was applied for the volume changes of Pp-side HA or Pp-opposite-side HA.

**Figure 7.**
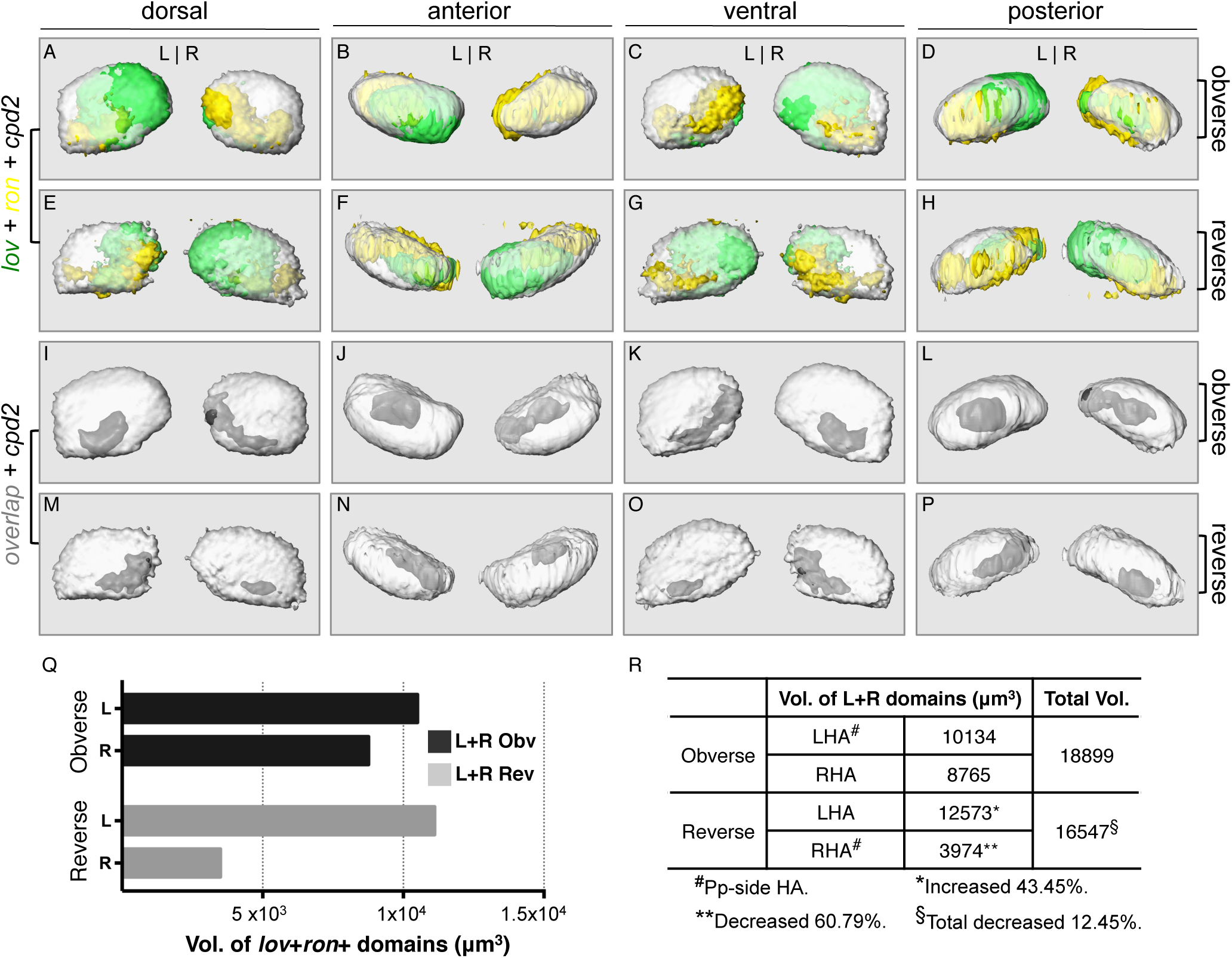
Decrease of *lov* and *ron* double-positive domains in left-right reversed habenular nuclei in 4 dpf larvae. (A-H) Snapshots from virtual 3D map from four different angles showing merged maps of *cpd2, lov* and *ron* transcripts from obverse (A-D) or reverse (E-H) groups. (I-P) Snapshots showing *lov* and *ron* overlapped domains (in gray) from obverse (I-L) or reverse (M-P) groups. (Q) Statistical chart showing the expression volumes of the *lov* and *ron* overlapped (L+R) domains in obverse (Obv) or reverse (Rev) model maps. (R). A table showing the computed volumes of L+N domains in the virtual 3D map. Total changes in percentage was calculated by using (Rev total – Obv total) / Obv total X 100. Same logic was applied for the volume changes of Pp-side HA or Pp-opposite-side HA.

## DISCUSSION

In this study, we utilized 2-color fluorescent RNA *in situ* hybridization (FISH) and the customized image processing protocol in Amira software to address the questions that previously were not addressed before (Figure 1). Due to the 3D nature of a developing brain, the adoption of FISH rather than 2D colorimetric gene expression analysis approach enabled us to uncover interesting new insights into the developmental differences between zebrafish larvae carrying different HA laterality. First, simply through 3D gene expression volume analyses, we found that the expression domain sizes of four commonly used developmental marker genes of HA (*cpd2, lov, nrp1a* and *ron*) in LHA and RHA were not all mirroringly reversed in HA laterality reversed larval brains (Figure 3I-M). These observations were further reflected by the analyses of L-R gene expression volume ratios for *cpd2, lov* and *ron* (Figure 3N), and thus proved that 2D snapshots of ISH results although may reflect certain degrees of gene expression alterations in many cases, it however cannot reflect the 3 dimensional gene expression changes within given populations of wildtype larvae carrying obverse or reversed brain laterality. We speculate that this claim may also be applicable to other cases where 2D snapshots are adopted for comparing gene expression domain sizes that are existed in 3D in experimental samples. One study in 2017 indicates that traditional behavioral scoring of individual zebrafish performed in 2D rather than 3D are flawed by over-reporting and under-reporting of locomotory differences (Macri et al., 2017). Therefore, to better represent the distribution of cells in neural tissues that inhere more differentiated cell types than any other tissues, spatial descriptions of gene expression patterns in 3D should be adopted because these neurons are organized in 3D in developing larvae.

To echo our study needs and the facts that 3D gene expression atlases are used progressively more to uncover new features of gene expression patterns (Akiyama-Oda and Oda, 2016; Keranen et al., 2006; Luengo Hendriks et al., 2006; Yang et al., 2019), a method that allows simultaneously quantifying and visualizing multiple gene expressions in developing HA has been developed in this study. Comparing to the sample processing procedures that detected multiple transcripts in whole-mount zebrafish embryos utilizing techniques involving the use of specific fluorophore-labeled RNA hairpins or specifically designed short probes containing specific pre-amplifier-interacting tails for each gene of interests (Choi et al., 2010; Gross-Thebing et al., 2014), our probe detecting procedures could easily be adopted by most laboratories utilizing standard probe generation and detection protocol. In addition, our method could accommodate more genes of interests without increasing probe detection difficulty. Further more, comparing to the previously reported method that adopted similar image aligning concept (Asadulina et al., 2012), our method is the first integrated protocol that can align and average multiple gene expression volumes in one processing and visualization platform. Another feature is that instead of adopting the method that arbitrarily selected the reference image stacks and artificially distorted image shapes for image registration purpose (Tay et al., 2011), we have created a more objective approach to select the reference image stack from a given experimental population and generate the volume map free from image distortion by performing landmark free reciprocal rigid registration and image correlation analyses. This approach avoided a larger image capture region for landmarks that was irrelevant to the neuronal clusters or tissues of study interests.

The unique usefulness of the 3D multi-gene expression maps of HA generated in this study is that they allowed us to better quantify the development of some unique groups of HA neurons such as the *lov*-*nrp1a* or *lov*-*ron* double-positive cells in larval brains with different laterality. As a result, we found significant increase of total *lov*-*nrp1a* (87.78%, figure 6) and significant decrease of total *lov-ron* (24%, figure 7) double-positive neurons in the L-R reversed HA in 4 dpf larvae. These observations indicate that Pp-guided laterality switch although did not alter the overall neuronal numbers in developing habenula (Figure 1I), cell fate alterations were accompanying the Pp-guided laterality switch. This result thus provided a cellular basis for the HA laterality-switch-associated behavior changes. In addition, because mirrored switch of gene expressions occurred to some but not all gene in HA with the Pp position switch, it explained why L-R reversals do not necessarily represent a neuroanatomical mirror image (Facchin et al., 2015).

In conclusion, generation of the 3D multi-gene expression volume maps provide us the tool to analyze the relative contribution of each key regulatory molecule on the development of the different neuronal lineages and to distinguish differences between normally and abnormally developed neuronal clusters in the embryonic HA. Additions of more gene markers into our map may help researchers to understand the complex differentiation process of the vertebrate HA. The image processing method we developed in this report also provides researchers a useful tool to establish multi-gene expression maps in 3D in the other tissues.

## ACKNOWLEDGMENTS

We thank Taiwan Zebrafish Core Facility funded by MOST (Ministry of Science and Technology, Taiwan) Grant #108-2319-B-400-002 for fish line maintenance and reagents. We also thank Drs. S. W. Wilson (UCL, UK), Sheng-Ping L. Hwang (ICOB, Academia Sinica, Taiwan) and Bon-Chu Chung (IMB, Academia Sinica) for critical discussion; Dr. M. E. Halpern (Carnegie Inst. for Science) for reagents; Dr. Sheng-Wei Lin (IBC, Academia Sinica, Taiwan) and Mr. Jaguar Li (ChingYeh Corp., Taiwan) for technical supports.

## AUTHOR CONTRIBUTIONS

G-T Wang, C-H Hsieh and Y-S Kuan designed the experiments. Y-S Kuan provided the reagents. G-T Wang, H-Y Pan, W-H Lang, Y-D Yu and Y-S Kuan performed the experiments. G-T Wang and Y-S Kuan analyzed the data. Y-S Kuan wrote the paper with the inputs from Jaguar Li and G-T Wang.

## SUPPLEMENTARY MATERIAL

The supplemental data for this article include one Supplementary Figure and two Supplementary Tables that can be downloaded at https://xxx.xxx.xxx.xxx (to be announced once published).

