## Supplementary Figures and Tables for "Three Dimensional Multi-gene Expression Maps Reveal Cell Fate Changes Associated with Laterality Reversal of Zebrafish Habenula"

**Supplementary Figure 1. Threshold and bleaching tests of fluorescent signals.** (A-D') Representative Z-projection images showing their original acquired signals (A-D) or their noise eliminated signals (A'-D'). I = intensity (Range is 0-255.). (E-H) Images showing Cy3 signals (*cpd2)* or FLU signals (*lov)* being collected the 1$^{st}$ time (E, G) or the 5$^{th}$ time. (I) Volumes from 2 Z-stacks that were under signal collection for 5 times (Z1-1 to Z1-5, or Z2-1 to Z2-5). Unit = μm$^3$. (H) Plots showing the volumes from (I).

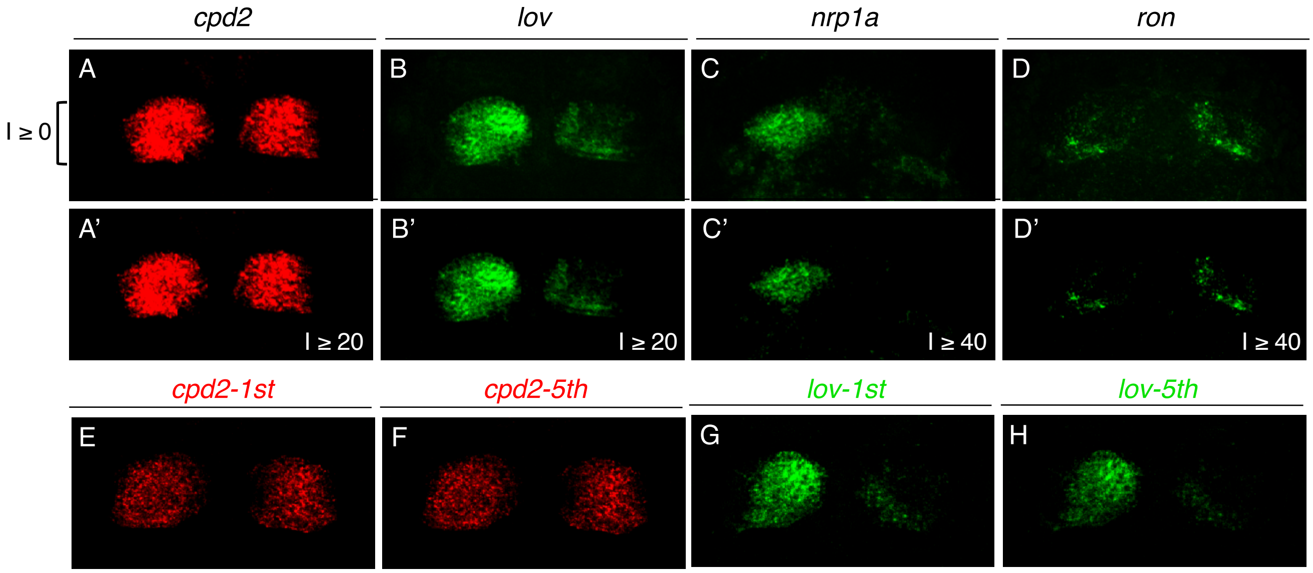


I

| | ***cpd2*-L** | ***cpd2*-R** | ***lov*-L** | ***lov*-R** |
|---|---|---|---|---|
| **Z1-1** | 91,611 | 80,222 | 101,079 | 24,194 |
| **Z1-2** | 87,023 | 76,353 | 88,279 | 17,065 |
| **Z1-3** | 86,587 | 76,020 | 82,393 | 14,543 |
| **Z1-4** | 87,235 | 76,340 | 84,952 | 16,003 |
| **Z1-5** | 89,561 | 78,757 | 77,310 | 12,739 |
| **Z2-1** | 70,094 | 51,491 | 86,023 | 166,49 |
| **Z2-2** | 70,420 | 51,855 | 79,738 | 134,99 |
| **Z2-3** | 73,308 | 54,199 | 69,975 | 10,331 |
| **Z2-4** | 69,437 | 51,352 | 62,116 | 7,635 |
| **Z2-5** | 69,917 | 51,724 | 58,693 | 6,544 |

J

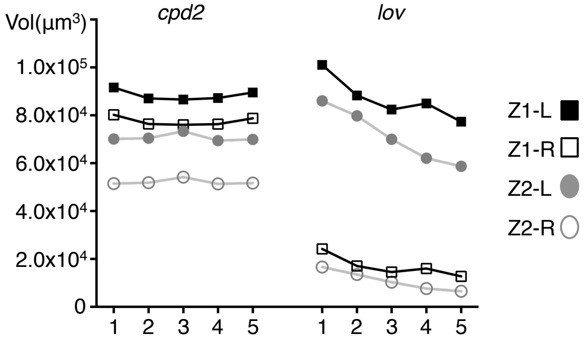

**Supplementary Table 1. Computed correlation factors from reciprocal one-to-one registration of each Z-stacks of *cpd2*.** (A) Correlation factors (CFs) from reciprocal one-to-one registeration between each Z-stacks of *cpd2* gene in obverse group. (B) CFs from samples of reversed group. Refer to Materials and Methods for details. LC1 to LC9 or rLC1 to rLC6 represent the 9 or 6 samples that were stained with *lov* (L) and *cpd2* (C) antisense probes in obverse or reversed group (respectively). The same naming logic applied to the samples that were stained with *nrp1a* (N) and *cpd2* (C), or with *ron* (R) and *cpd2* (C) antisense probes.

A.

| Name | LC1 | LC2 | LC3 | LC4 | LC5 | LC6 | LC7 | LC8 | LC9 | NC1 | NC2 | NC3 | NC4 | NC5 | NC6 | RC1 | RC2 | RC3 | RC4 | RC5 | RC6 | RC7 | RC8 |
|---|---|---|---|---|---|---|---|---|---|---|---|---|---|---|---|---|---|---|---|---|---|---|---|
| LC1 | 1 | 0.82 | 0.82 | 0.77 | 0.77 | 0.88 | 0.81 | 0.82 | 0.83 | 0.85 | 0.64 | 0.68 | 0.74 | 0.82 | 0.74 | 0.78 | 0.81 | 0.83 | 0.84 | 0.78 | 0.81 | 0.86 | 0.83 |
| LC2 | 0.82 | 1 | 0.28 | 0.76 | 0.81 | 0.86 | 0.78 | 0.83 | 0.79 | 0.80 | 0.60 | 0.66 | 0.72 | 0.78 | 0.71 | 0.72 | 0.83 | 0.86 | 0.85 | 0.78 | 0.80 | 0.85 | 0.85 |
| LC3 | 0.82 | 0.78 | 1 | 0.78 | 0.76 | 0.82 | 0.79 | 0.80 | 0.83 | 0.83 | 0.63 | 0.70 | 0.73 | 0.82 | 0.77 | 0.77 | 0.78 | 0.82 | 0.82 | 0.78 | 0.81 | 0.84 | 0.82 |
| LC4 | 0.77 | 0.76 | 0.78 | 1 | 0.76 | 0.77 | 0.77 | 0.77 | 0.79 | 0.77 | 0.67 | 0.70 | 0.73 | 0.77 | 0.75 | 0.75 | 0.73 | 0.80 | 0.77 | 0.72 | 0.74 | 0.79 | 0.77 |
| LC5 | 0.77 | 0.81 | 0.76 | 0.76 | 1 | 0.80 | 0.77 | 0.82 | 0.77 | 0.79 | 0.67 | 0.62 | 0.75 | 0.78 | 0.73 | 0.74 | 0.80 | 0.82 | 0.81 | 0.75 | 0.77 | 0.81 | 0.80 |
| LC6 | 0.88 | 0.86 | 0.82 | 0.77 | 0.79 | 1 | 0.81 | 0.84 | 0.83 | 0.85 | 0.61 | 0.68 | 0.73 | 0.81 | 0.73 | 0.76 | 0.83 | 0.87 | 0.88 | 0.81 | 0.83 | 0.88 | 0.86 |
| LC7 | 0.81 | 0.78 | 0.79 | 0.77 | 0.77 | 0.81 | 1 | 0.80 | 0.81 | 0.81 | 0.65 | 0.68 | 0.72 | 0.79 | 0.74 | 0.75 | 0.77 | 0.81 | 0.80 | 0.75 | 0.76 | 0.82 | 0.79 |
| LC8 | 0.82 | 0.83 | 0.81 | 0.77 | 0.82 | 0.84 | 0.80 | 1 | 0.82 | 0.83 | 0.64 | 0.69 | 0.75 | 0.81 | 0.74 | 0.74 | 0.83 | 0.86 | 0.85 | 0.80 | 0.81 | 0.85 | 0.85 |
| LC9 | 0.83 | 0.79 | 0.83 | 0.78 | 0.77 | 0.83 | 0.80 | 0.82 | 1 | 0.85 | 0.66 | 0.71 | 0.74 | 0.81 | 0.77 | 0.75 | 0.80 | 0.84 | 0.82 | 0.79 | 0.81 | 0.84 | 0.83 |
| NC1 | 0.85 | 0.80 | 0.83 | 0.77 | 0.78 | 0.85 | 0.81 | 0.83 | 0.84 | 1 | 0.66 | 0.72 | 0.76 | 0.85 | 0.78 | 0.79 | 0.83 | 0.84 | 0.87 | 0.83 | 0.84 | 0.86 | 0.86 |
| NC2 | 0.64 | 0.61 | 0.65 | 0.67 | 0.67 | 0.62 | 0.66 | 0.64 | 0.66 | 0.67 | 1 | 0.68 | 0.70 | 0.70 | 0.71 | 0.71 | 0.62 | 0.64 | 0.63 | 0.62 | 0.64 | 0.66 | 0.63 |
| NC3 | 0.69 | 0.68 | 0.71 | 0.71 | 0.72 | 0.69 | 0.68 | 0.70 | 0.72 | 0.73 | 0.68 | 1 | 0.74 | 0.76 | 0.73 | 0.73 | 0.71 | 0.72 | 0.71 | 0.72 | 0.71 | 0.72 | 0.71 |
| NC4 | 0.74 | 0.72 | 0.73 | 0.72 | 0.75 | 0.73 | 0.72 | 0.75 | 0.74 | 0.76 | 0.69 | 0.74 | 1 | 0.77 | 0.75 | 0.76 | 0.73 | 0.75 | 0.75 | 0.74 | 0.76 | 0.75 | 0.75 |
| NC5 | 0.82 | 0.78 | 0.82 | 0.75 | 0.77 | 0.81 | 0.78 | 0.81 | 0.81 | 0.85 | 0.69 | 0.75 | 0.77 | 1 | 0.8 | 0.82 | 0.83 | 0.82 | 0.84 | 0.84 | 0.82 | 0.83 | 0.84 |
| NC6 | 0.74 | 0.71 | 0.77 | 0.74 | 0.73 | 0.73 | 0.74 | 0.74 | 0.77 | 0.78 | 0.71 | 0.72 | 0.75 | 0.80 | 1 | 0.79 | 0.72 | 0.75 | 0.76 | 0.75 | 0.75 | 0.76 | 0.75 |
| RC1 | 0.78 | 0.72 | 0.76 | 0.74 | 0.74 | 0.76 | 0.75 | 0.74 | 0.75 | 0.79 | 0.70 | 0.71 | 0.76 | 0.82 | 0.78 | 1 | 0.72 | 0.74 | 0.78 | 0.77 | 0.78 | 0.77 | 0.76 |
| RC2 | 0.82 | 0.85 | 0.78 | 0.74 | 0.78 | 0.86 | 0.77 | 0.84 | 0.80 | 0.84 | 0.58 | 0.70 | 0.73 | 0.83 | 0.72 | 0.75 | 1 | 0.87 | 0.90 | 0.88 | 0.85 | 0.87 | 0.90 |
| RC3 | 0.83 | 0.86 | 0.82 | 0.80 | 0.82 | 0.87 | 0.80 | 0.86 | 0.84 | 0.85 | 0.63 | 0.71 | 0.75 | 0.82 | 0.74 | 0.75 | 0.86 | 1 | 0.88 | 0.82 | 0.85 | 0.88 | 0.89 |
| RC4 | 0.84 | 0.85 | 0.82 | 0.77 | 0.80 | 0.88 | 0.79 | 0.84 | 0.82 | 0.88 | 0.62 | 0.71 | 0.75 | 0.85 | 0.76 | 0.78 | 0.89 | 0.88 | 1 | 0.86 | 0.86 | 0.88 | 0.90 |
| RC5 | 0.80 | 0.77 | 0.78 | 0.72 | 0.73 | 0.81 | 0.75 | 0.80 | 0.79 | 0.84 | 0.58 | 0.71 | 0.74 | 0.84 | 0.72 | 0.77 | 0.88 | 0.82 | 0.86 | 1 | 0.86 | 0.83 | 0.87 |
| RC6 | 0.81 | 0.80 | 0.81 | 0.74 | 0.77 | 0.83 | 0.76 | 0.81 | 0.81 | 0.84 | 0.62 | 0.70 | 0.75 | 0.82 | 0.75 | 0.78 | 0.85 | 0.85 | 0.86 | 0.86 | 1 | 0.84 | 0.87 |
| RC7 | 0.86 | 0.85 | 0.83 | 0.77 | 0.81 | 0.89 | 0.82 | 0.85 | 0.84 | 0.86 | 0.65 | 0.70 | 0.75 | 0.83 | 0.76 | 0.77 | 0.86 | 0.88 | 0.88 | 0.83 | 0.84 | 1 | 0.88 |
| RC8 | 0.83 | 0.85 | 0.82 | 0.76 | 0.80 | 0.87 | 0.78 | 0.85 | 0.83 | 0.86 | 0.62 | 0.70 | 0.75 | 0.84 | 0.75 | 0.76 | 0.90 | 0.89 | 0.90 | 0.87 | 0.87 | 0.88 | 1 |
| Ave. | 0.81 | 0.79 | 0.80 | 0.76 | 0.78 | 0.82 | 0.78 | 0.81 | 0.80 | 0.82 | 0.66 | 0.71 | 0.75 | 0.81 | 0.76 | 0.77 | 0.81 | 0.82 | 0.83 | 0.80 | 0.81 | 0.83 | 0.83 |
| STD | 0.07 | 0.08 | 0.06 | 0.06 | 0.06 | 0.08 | 0.06 | 0.07 | 0.06 | 0.06 | 0.08 | 0.07 | 0.06 | 0.05 | 0.06 | 0.06 | 0.08 | 0.07 | 0.07 | 0.07 | 0.07 | 0.07 | 0.08 |

B.

| Name | rLC1 | rLC2 | rLC3 | rLC4 | rLC5 | rLC6 | rNC1 | rNC2 | rNC3 | rNC4 | rRC1 | rRC2 | rRC3 | rRC4 |
|---|---|---|---|---|---|---|---|---|---|---|---|---|---|---|
| rLC1 | 1 | 0.74 | 0.77 | 0.63 | 0.73 | 0.76 | 0.59 | 0.75 | 0.77 | 0.36 | 0.76 | 0.77 | 0.69 | 0.79 |
| rLC2 | 0.74 | 1 | 0.70 | 0.75 | 0.79 | 0.79 | 0.70 | 0.76 | 0.81 | 0.46 | 0.85 | 0.82 | 0.84 | 0.80 |
| rLC3 | 0.76 | 0.70 | 1 | 0.54 | 0.71 | 0.78 | 0.63 | 0.72 | 0.77 | 0.56 | 0.75 | 0.70 | 0.65 | 0.80 |
| rLC4 | 0.63 | 0.75 | 0.53 | 1 | 0.70 | 0.62 | 0.74 | 0.65 | 0.68 | 0.69 | 0.75 | 0.71 | 0.81 | 0.69 |
| rLC5 | 0.73 | 0.79 | 0.71 | 0.71 | 1 | 0.76 | 0.64 | 0.73 | 0.76 | 0.64 | 0.81 | 0.78 | 0.78 | 0.80 |
| rLC6 | 0.76 | 0.79 | 0.78 | 0.71 | 0.78 | 1 | 0.66 | 0.74 | 0.80 | 0.69 | 0.83 | 0.79 | 0.81 | 0.83 |
| rNC1 | 0.51 | 0.70 | 0.65 | 0.73 | 0.62 | 0.64 | 1 | 0.51 | 0.59 | 0.46 | 0.69 | 0.65 | 0.79 | 0.58 |
| rNC2 | 0.74 | 0.76 | 0.72 | 0.66 | 0.73 | 0.73 | 0.57 | 1 | 0.80 | 0.58 | 0.77 | 0.77 | 0.72 | 0.79 |
| rNC3 | 0.76 | 0.81 | 0.77 | 0.69 | 0.78 | 0.78 | 0.59 | 0..80 | 1 | 0.64 | 0.85 | 0.80 | 0.76 | 0.86 |
| rNC4 | 0.43 | 0.68 | 0.49 | 0.68 | 0.63 | 0.60 | 0.49 | 0.57 | 0.62 | 1 | 0.44 | 0.67 | 0.73 | 0.62 |
| rRC1 | 0.76 | 0.85 | 0.75 | 0.76 | 0.82 | 0.78 | 0.70 | 0.78 | 0.86 | 0.41 | 1 | 0.82 | 0.85 | 0.88 |
| rRC2 | 0.77 | 0.82 | 0.70 | 0.71 | 0.78 | 0.78 | 0.64 | 0.77 | 0.80 | 0.68 | 0.82 | 1 | 0.81 | 0.82 |
| rRC3 | 0.69 | 0.84 | 0.61 | 0.81 | 0.79 | 0.74 | 0.79 | 0.72 | 0.77. | 0.75 | 0.85 | 0.81 | 1 | 0.79 |
| rRC4 | 0.79 | 0.82 | 0.81 | 0.71 | 0.81 | 0.83 | 0.52 | 0.80 | 0.87 | 0.67 | 0.89 | 0.83 | 0.81 | 1 |
| Ave. | 0.72 | 0.79 | 0.71 | 0.72 | 0.76 | 0.76 | 0.66 | 0.73 | 0.78 | 0.61 | 0.79 | 0.78 | 0.79 | 0.79 |
| STD | 0.13 | 0.08 | 0.13 | 0.10 | 0.09 | 0.10 | 0.13 | 0.11 | 0.10 | 0.16 | 0.13 | 0.09 | 0.08 | 0.10 |

**Supplementary Table 2. Expression domain volumes and their left-to-right (L/R) ratios.** LC1-9, NC1-6 and RC1-8 for obverse group; rLC1-6, rNC1-4 and rRC1-4 for reversed group. LC represents samples stained with *lov* + *cpd2* antisense probes. LN or LR are for *cpd2* + *nrp1a* antisense probes or *cpd2* + *ron* antisense probes .

| *cpd2* | Left ($\mu m^3$) | Right ($\mu m^3$) | L/R % | *cpd2* | Left ($\mu m^3$) | Right ($\mu m^3$) | L/R % |
|---|---|---|---|---|---|---|---|
| **LC1** | 123,136 | 98,368 | 125 | **rLC1** | 56,116 | 71,252 | 79 |
| **LC2** | 111,824 | 79,198 | 141 | **rLC2** | 77,593 | 83,875 | 93 |
| **LC3** | 87,881 | 80,772 | 109 | **rLC3** | 96,735 | 111,244 | 87 |
| **LC4** | 68,849 | 51,925 | 133 | **rLC4** | 63,608 | 74,010 | 86 |
| **LC5** | 75,775 | 52,624 | 144 | **rLC5** | 62,434 | 67,980 | 92 |
| **LC6** | 136,127 | 92,703 | 147 | **rLC6** | 79,594 | 115,984 | 67 |
| **LC7** | 86,100 | 69,734 | 123 | | | | |
| **LC8** | 94,747 | 70,168 | 135 | | | | |
| **LC9** | 988,13 | 84,454 | 117 | | | | |
| **NC1** | 119,622 | 91,248 | 131 | **rNC1** | 89,079 | 93,526 | 95 |
| **NC2** | 41,901 | 29,747 | 141 | **rNC2** | 64,889 | 71,149 | 91 |
| **NC3** | 51,471 | 35,943 | 143 | **rNC3** | 99,605 | 114,113 | 87 |
| **NC4** | 69,043 | 44,135 | 156 | **rNC4** | 115,332 | 133,189 | 87 |
| **NC5** | 96,760 | 88,909 | 109 | | | | |
| **NC6** | 67,717 | 54,434 | 124 | | | | |
| **RC1** | 86,525 | 55,635 | 156 | **rRC1** | 96,385 | 112,305 | 86 |
| **RC2** | 126,262 | 89,212 | 142 | **rRC2** | 79,970 | 81,938 | 98 |
| **RC3** | 116,637 | 81,951 | 142 | **rRC3** | 96,466 | 96,573 | 99 |
| **RC4** | 128,221 | 93,959 | 136 | **rRC4** | 91,538 | 111,315 | 82 |
| **RC5** | 122,432 | 87,223 | 140 | | | | |
| **RC6** | 131,815 | 88,144 | 150 | | | | |
| **RC7** | 132,484 | 95,458 | 139 | | | | |
| **RC8** | 136,210 | 102,731 | 133 | | | | |
| **Ave.** | 100,450 | 74,725 | 136 | **Ave.** | 83,524 | 95,604 | 88 |
| **STD** | 28,488 | 21,302 | 13 | **STD** | 17,267 | 20,923 | 8 |

| *lov* | Left ($\mu m^3$) | Right ($\mu m^3$) | L/R % | *lov* | Left ($\mu m^3$) | Right ($\mu m^3$) | L/R % |
|---|---|---|---|---|---|---|---|
| **LC1** | 79,766 | 7,118, | 1,121 | **rLC1** | 86,262 | 136,997 | 63 |
| **LC2** | 70,937 | 9,846 | 720 | **rLC2** | 46,301 | 106,713 | 43 |
| **LC3** | 68,910 | 13,025 | 529 | **rLC3** | 78,131 | 125,921 | 62 |
| **LC4** | 42,107 | 5,402 | 779 | **rLC4** | 63,392 | 73,945 | 86 |
| **LC5** | 48,877 | 3,938 | 1,241 | **rLC5** | 29,691 | 70,374 | 42 |
| **LC6** | 94,956 | 14,843 | 640 | **rLC6** | 47,363 | 117,814 | 40 |
| **LC7** | 86,182 | 51,494 | 167 | | | | |
| **LC8** | 117,841 | 39,737 | 297 | | | | |
| **LC9** | 117,939 | 31,062 | 380 | | | | |
| **Ave.** | 80,835 | 19,607 | 653 | **Ave.** | 58,523 | 102,790 | 56 |
| **STD** | 26,778 | 17,021 | 369 | **STD** | 21,368 | 27,537 | 28 |

| *nrp1a* | Left ($\mu m^3$) | Right ($\mu m^3$) | L/R % | *nrp1a* | Left ($\mu m^3$) | Right ($\mu m^3$) | L/R % |
|---|---|---|---|---|---|---|---|
| **NC1** | 44,581 | 51 | N/A | **rNC1** | 8,672 | 66,566 | 13 |
| **NC2** | 17,816 | 69 | N/A | **rNC2** | 12,511 | 44,699 | 28 |
| **NC3** | 28,744 | 180 | N/A | **rNC3** | 31,345 | 41,228 | 76 |
| **NC4** | 17,935 | 70 | N/A | **rNC4** | 16,770 | 67,426 | 25 |
| **NC5** | 62,120 | 3,990 | N/A | | | | |
| **NC6** | 34,623 | 234 | N/A | | | | |
| **Ave.** | 34,303 | 766 | | **Ave.** | 17,324 | 54,980 | 35 |
| **STD** | 17,030 | 1,581 | | **STD** | 9,915 | 13,952 | 28 |

| *ron* | Left ($\mu m^3$) | Right ($\mu m^3$) | L/R % | *ron* | Left ($\mu m^3$) | Right ($\mu m^3$) | L/R % |
|---|---|---|---|---|---|---|---|
| **RC1** | 6,581 | 11,525 | 57 | **rRC1** | 28,125 | 9,957 | 282 |
| **RC2** | 16,667 | 40,051 | 42 | **rRC2** | 8.663 | 3,524 | 246 |
| **RC3** | 34,274 | 37,487 | 91 | **rRC3** | 15,275 | 7,589 | 201 |
| **RC4** | 22,349 | 31,929 | 70 | **rRC4** | 13,789 | 3,269 | 422 |
| **RC5** | 32,133 | 47,789 | 67 | | | | |
| **RC6** | 28, 799 | 38,467 | 75 | | | | |
| **RC7** | 11,149 | 23,635 | 47 | | | | |
| **RC8** | 9,986 | 16,511 | 60 | | | | |
| **Ave.** | 20,242 | 30,924 | 64 | **Ave.** | 16,463 | 6,085 | 288 |
| **STD** | 10,712 | 12,569 | 15 | **STD** | 8,275 | 3,253 | 95 |
